# Modeling Nitric Oxide Diffusion and Plasticity Modulation in Cerebellar Learning

**DOI:** 10.1101/2024.07.24.604957

**Authors:** Alessandra Maria Trapani, Carlo Andrea Sartori, Benedetta Gambosi, Alessandra Pedrocchi, Alberto Antonietti

## Abstract

Nitric Oxide (NO) is a versatile signalling molecule with significant roles in various physiological processes, including synaptic plasticity and memory formation. In the cerebellum, NO is produced by neural NO Synthase and diffuses to influence synaptic changes, particularly at parallel fiber - Purkinje cell synapses. This study aims to investigate NO’s role in cerebellar learning mechanisms using a biologically realistic simulation-based approach. We developed the NO Diffusion Simulator (NODS), a Python module designed to model NO production and diffusion within a cerebellar spiking neural network framework. Our simulations focus on the Eye-Blink Classical Conditioning protocol to assess the impact of NO modulation on long-term potentiation and depression at parallel fiber - Purkinje cell synapses. The results demonstrate that NO diffusion significantly affects synaptic plasticity, dynamically adjusting learning rates based on synaptic activity patterns. This metaplasticity mechanism enhances the cerebellum’s capacity to prioritize relevant inputs and mitigate learning interference selectively modulating synaptic efficacy. Our findings align with theoretical models suggesting that NO serves as a contextual indicator, optimizing learning rates for effective motor control and adaptation to new tasks. The NODS implementation provides an efficient tool for large-scale simulations, facilitating future studies on NO dynamics in various brain regions and neurovascular coupling scenarios. By bridging the gap between molecular processes and network-level learning, this work underscores the critical role of NO in cerebellar function and offers a robust framework for exploring NO-dependent plasticity in computational neuroscience.

## 1 Introduction

Nitric Oxide (NO) is an essential molecule ubiquitously present in the human organism with a significant role in various physiological processes, spacing from muscular relaxation to immune response, from vascular dilatation to neurovascular coupling, and coordination of long-term memory [Koshland, 1992]. Over the past 35 years, extensive research has been conducted to explore the diverse pathways involved in NO signalling, its diffusion mechanisms, and its impact on synaptic transmission. This research has provided valuable insights into the functional consequences of NO in both normal physiological and pathological states.

What makes NO particularly intriguing as a signalling molecule is that, unlike other neurotransmitters that act on specific receptors expressed in dedicated neurons, NO is a versatile signal that can influence a wide range of neurons through diverse molecular mechanisms. The low molecular weight of NO and its diffusion coefficient in aqueous environment confer to it the possibility to uniformly disperse in all directions from its site of production, enabling forms of cellular communication that do not rely just on anatomical connections of synaptic boutons [Garthwaite, 2016]. NO in the Central Nervous System is produced by the enzyme *neural NO Synthase* (nNOS), which is expressed in specific neuronal subtypes and can be located in the cytoplasm for small subpopulations of GABAergic cells or in the spine head for larger populations of excitatory neurons. nNOS is physically tethered to NMDA receptors (NMDAr), which are fundamental gates for *Ca*^2+^ signalling, and it starts producing NO after the [*Ca*^2+^] increase, following the activation of NMDAr by glutamate [Brenman et al., 1996, Garthwaite, 2016]. Following production and diffusion, at given concentrations NO activates different possible biochemical cascades. Inside cells it reacts with soluble Guanylyl Cyclase (sGC), its most sensitive target found in the organism, or in an sGC-independent manner by modification of the thiol (-SH) side chains of cysteine residues in proteins [Garthwaite, 2016]. Experimentally, [*NO*] has been measured in a range that spans tens of pM to nM to be sufficient to activate the NO signalling cascade [Garthwaite et al., 1988, Hardingham et al., 2013]. Furthermore, researchers tried to estimate the inactivation rate for the NO in brain tissue and its maximum diffusional range, by directly measuring NO concentration or by modelling its kinetics to account for experimental measures of other molecules present in its biochemical cascade. Laranjinha et al. [2012] attempted it by employing NO-selective carbon fibre microelectrodes to directly measure alterations in NO levels following the localized administration of small volumes *in vivo*. In early modelling examples to characterize the volumetric signalling properties of NO [Ning Feng et al., 2005, Smith and Philippides, 2000], a decay rate value of *λ* = 0.1386 *s*^−1^ (corresponding to a half-life of 5 s) was commonly chosen. However, for scenarios involving strong NO sinks, Philippides et al. [2000] used an inactivation rate of *λ* = 693.15 *s*^−1^ (equivalent to a half-life of 1 ms), representing a conservative estimate based on the rate of NO uptake by nearby haemoglobin-containing structures like blood vessels. Garthwaite [2016] modelled the source structure (nNOS) and surroundings as a disc of diameter around 400 nm that constantly produces NO with a rate of 40 *molecule/s*^−1^. With a single synaptic stimulus, the peak value of [NO], after 40 ms, is 60 pM at the center of the source and steeply decreases to 5 pM 1*μ*m away from the center.

### 1.1 NO as second messenger in the brain

Extensive research has been conducted to explore the effects of NO in many brain areas, such as the visual cortex [Haghikia et al., 2007, Liu and Zweier, 2013, Volgushev et al., 2000], where NO signalling has been identified as a crucial factor in the induction of long-term potentiation (LTP) [Haghikia et al., 2007, Liu and Zweier, 2013, Volgushev et al., 2000]. The findings on the involvement of NO in cortical LTP are further reinforced by the impaired neocortical LTP observed in eNOS-deficient mice [Haul et al., 1999], as well as the identification of a NO-dependent component of LTP in the barrel cortex [Hardingham and Fox, 2006, Dachtler et al., 2012], medial frontal cortex [Nowicky and Bindman, 1993], and rat auditory cortex [Wakatsuki et al., 1998]. Furthermore, in the hippocampus, NO has been commonly observed to participate in later phases of LTP, whereby inhibition of NO synthase results in a gradual decay back towards the baseline while leaving the early potentiation largely unaffected [Phillips et al., 2008], while in the thalamus increasing NO levels resulted in the potentiation of inhibitory activity, likely through a cGMP-dependent process that enhances GABA release from presynaptic terminals [Yang and Cox, 2007]. The changes in ion channel function caused by NO signalling impact neuronal excitability, synaptic transmission, and the propagation of action potentials. In the past few years, an increasing number of studies (reviewed in Hardingham et al. [2013]) suggested that certain stimulation patterns of a closely packed group of neurons, containing neuronal nNOS enzyme, may generate a diffuse cloud of NO, thus acting as a volume transmitter, with a relatively large area of influence. On the other hand, isolated stimuli would just lead to a local effect of the NO signal, exerting classical communication through a single anatomical synaptic connection [Garthwaite, 2016].

### 1.2 NO physiological role in the cerebellum

Many research studies found proof for the presence of nNOS in the cerebellum both at the Granular Layer and Molecular Layer. These were followed by others that covered various aspects ranging from the NO signalling cascade to its impact on synapses and even behavioural consequences associated with deficiencies or excessive levels of NO [Lev-Ram et al., 2002, Maffei et al., 2003, Jacoby et al., 2001, Reynolds and Hartell, 2001, Wang et al., 2014, Namiki et al., 2005, Bouvier et al., 2016, Kimura et al., 1998]. The authors observed cerebellar slices where nNOS is expressed in Granule Cells (GrC) and Molecular Interneurons [Lev-Ram et al., 2002, Wang et al., 2014, Kono et al., 2019].

A high number of nNOS was found also in the Purkinje Cell (PC) dendritic tree, meaning that the NO production happens in GrCs, both in the granular layer at the mossy fibers (*mf* s) - GrC synapses and at parallel fibers (*pf* s) -PC synapses [Wood et al., 2011]. Regarding synaptic plasticity, LTP induction at the level of *mf* -GrCs synapses relies on the activation of postsynaptic NMDA receptors, and it results blocked when inhibiting various steps of the NO cascade, including NOS-mediated NO production, NO diffusion in the extracellular space, and sGC activation. [Maffei et al., 2003]. In the molecular layer, many experimental studies investigated the role of NO in modulating synaptic plasticity at the *pf* -PC synapses [Moncada et al., 1989, Wood et al., 2011, Katoh et al., 2000]. Although it is established that NO is produced pre-synaptically, there is still ongoing debate regarding where in the synapse and how the effects of NO are exerted. Lev-Ram and colleagues conducted two separate studies demonstrating the necessity of NO in both long-term depression (LTD) and LTP at *pf* -PC synapses, with the two differing for the stimulus frequency needed at the *pf* s and the biochemical cascade exerted by NO [Lev-Ram et al., 2002, 1997]. Moreover, a more recent study reported a form of NO-dependent LTP, that similarly to LTD, is induced by high-frequency bursts of activity from the *pf* s [Bouvier et al., 2016]. While with 1 Hz stimulation LTP required the simultaneous involvement of presynaptic and postsynaptic activities [Wang et al., 2014]. Furthermore, in Jacoby et al. [2001], similarly to Reynolds and Hartell [2001], they focused on investigating the spread of potentiation to non-stimulated synapses and the underlying mechanisms following 8 Hz stimuli at *pf* s. Their findings suggest that heterosynaptic potentiation is pre-synaptically initiated. NO acts as a diffusible messenger facilitating presynaptic transmitter release within a radius of at least 150 *μ*m and this is in line with the results from Lev-Ram et al. [2002]. According to the latter, these findings suggest that the NO-dependent LTP may function as a plausible anti-Hebbian mechanism counterbalancing the effects of LTD and facilitating the reversal of motor learning.

Altogether, these research studies evidence an important role for NO in the cerebellum characterized by a diffusive plasticity effect, beyond the single synaptic connection.

### 1.3 Modelling NO diffusive plasticity in the cerebellum

Understanding the role of NO in neural processes is crucial for unravelling the mechanisms underlying brain learning and information processing. However, studying NO’s function *in vivo* and *in vitro* is challenging due to the difficulties in implementing experiments and measuring its highly volatile nature.

To address these challenges and gain deeper insights into NO’s role, a simulation-based approach becomes indispensable. By utilizing *in silico* simulations, we can overcome the limitations of traditional experimental methods and investigate the impact of NO in a more repeatable and controlled environment.

Abraham and Bear proposed the term *“metaplasticity”* to describe the influences that the prior history of synaptic activity has on the synaptic state, and thereby the degree of LTP or LTD produced by a given experimental protocol [Abraham and Bear, 1996]. Metaplasticity is set to prevent previously potentiated (depressed) synapses from undergoing further LTP (LTD), avoiding a saturated state. A second important function of metaplasticity may be to integrate synaptic events over much longer periods than the tens of milliseconds typical of temporal summation of synaptic potentials. Schweighofer and Arbib proposed to integrate the learning rule that describes the plasticity at *pf* -PC synapses with a model for adapting the learning rate in an activity-dependent manner [Schweighofer and Ferriol, 2000]. In addition, they pose interesting theoretical considerations to support the necessity of including metaplasticity in the computational models for simulating cerebellar learning as it could have a role in:

1. **Control learning rates of *pf* -PC synapses** by tuning the synapse parameters to find optimal learning rates;
2. **Address the overfitting Problem** by concentrating weight updates in PC synapses whose adaptation is most relevant to learning an overall pattern, and thus addressing a possible “overfitting” problem given by the high number of *pf* s-PC synapses
3. **Give relevance to inputs and learning interference** by increasing the signal-to-noise ratio during learning tasks by minimizing the adaptation to irrelevant stimuli while increasing learning rates of inputs likely to be relevant.

However, metaplasticity was appointed as a role for NO only in a more recent study [Safaryan et al., 2017], where it was proposed as an appropriate candidate for implementing a type of plasticity that is non-specific and outperforms the standard plasticity model of *pf* -PC synapses. The model is based on the experimental evidence that plasticity requires the action of a diffusive and volatile second messenger able to induce plasticity in neighbouring non-active synapses, such as the NO. Furthermore, a simulation study that integrated established models of PC electrophysiology, calcium dynamics, and signalling pathways for *pf* -PC LTD [Ogasawara et al., 2007] highlighted the critical influence of local NO concentration in the induction of LTD and its impact on input specificity. Starting from the experimental evidence that the coincidence of pre- and postsynaptic firing alone is insufficient for LTD to occur and adequate levels of NO generated by the activation of surrounding *pfs* are necessary [Lev-Ram et al., 2002, Wang et al., 2014], Ogasawara and colleagues found that excessive activity in neighbouring *pfs* leads to NO accumulation, resulting in LTD in synapses that were not directly stimulated, therefore compromising input specificity. Based on these findings, they propose the hypothesis that NO serves as a contextual indicator as long as a small number of *pfs* codes for a movement. Thus, they predict sparse *pf* activity *in vivo* to preserve input specificity, as excessive activity would diminish the specificity of the learning process. In this scenario, the NO facilitates the selection and updating of internal models according to the relevance of a given context.

In this work, we aim to investigate the role of NO in cerebellar learning mechanisms, by employing a biologically realistic simulation-based approach. With respect to previous work, we propose a more comprehensive simulation study that incorporates NO production and diffusion model within a physiologically-inspired cerebellar structure, leveraging the detailed morphological and topological network [De Schepper et al., 2022]. We simulate a large-scale cerebellar spiking neural network (SNN) that can learn the Eye-Blink Classical Conditioning (EBCC) [Geminiani et al., 2022] and we show which are the effects when we activate NO modulation for LTP and LTD at *pf* -PC synapses.

## 2 Materials and methods

### 2.1 NODS: Nitric Oxide Diffusion Simulator

#### 2.1.1 Simulator structure

We implemented a Python module to simulate the NO diffusion inside a neural network that is compatible with different simulation environments, such as NEST [Gewaltig and Diesmann, 2007] and Brain [Stimberg et al., 2019], and more generally with any neuronal model written in Python. The code is available at https://github.com/alessandratrapani/NODS. The simulator takes as input the spatial coordinates ([*x, y, z*]) of the nNOS, the activity of the neurons that activate the nNOS (presynaptic spikes), and the spatial coordinates of the points where the user wants to evaluate the NO concentration ([*NO*]). The simulator’s output is the temporal profile of the NO signal in the evaluation points and (optional) the temporal and spatial profile of the NO diffused from each nNOS. The simulator structure can be divided into three main blocks:

1. **Geometry initialization**: we compute the relative distances between all the nNOS placed in the network and all the points where the user wants to evaluate the [*NO*]. Then, we link each evaluation point to the nearest nNOS within 15 *μ*m. This constraint has been chosen in accordance with Garthwaite [2016]. In this way, we can reduce the computational load at the level of the single source by restricting the diffusion within 15 *μ*m radius from the source.
2. **Single source computation**: for each nNOS, we simulate the spatial and temporal profile of the NO generated and diffused. The equations for the single source model are reported in subsubsection 2.1.2.
3. **Diffusion evaluation**: spatial and temporal summation of the [*NO*] provided by all nNOS surrounding each evaluation point, with respect to their relative distances, already computed in the initialization phase (first block).

NODS allows us to perform both offline simulations, where the network activity is loaded before simulating the NO production and diffusion, and online simulations, where the network activity and the NO diffusion run in parallel. In the latter configuration, for each *dt* = 1 ms of the simulated network activity, we compute the amount of NO diffused in *dt*.

#### 2.1.2 Single source model

##### Production equations

The dynamic of NO production depends on complex biochemical reaction cascade^1^ [Hall and Garthwaite, 2009]. We split the reaction cascade into two parts and represented them with two differential equations: we designed Equation 1 for describing the Ca^2+^/calmodulin binding, and Equation 2 to describe the activation of nNOS enzyme.

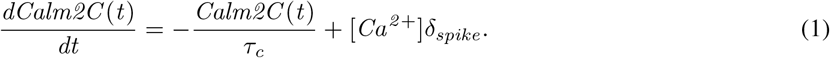

Where *τ*_*c*_ is a time constant describing the decay of Calm2C concentration after a spike of Ca^2+^ has entered the cell, [*Ca*^2+^]*δ*_*spike*_ in the equation. We chose *τ*_*c*_ = 150 ms and [*Ca*^2+^] = 1 [Sweeney et al., 2015].

The activation of the nNOS enzyme is modeled using the following equation:

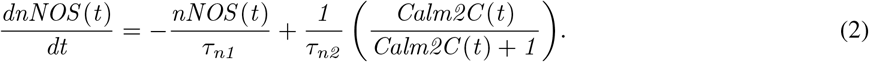

Where: *τ*_*n*1_ = 25 ms and *τ*_*n*2_ = 200 ms [Sweeney et al., 2015]. Here, we assumed that the amount of NO produced by a source is proportional to the amount of activated nNOS, given by Equation 2.

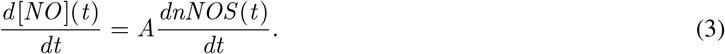

Where *A* = 1.35 10^−9^, better replicate the equivalent simulations performed with the simulator NEURON [Kumbhar et al., 2019] used as reference.

##### Diffusion equation

To model NO diffusion, we used the heat diffusion equation, where the solution is the NO concentration [*NO*](**x**, *t*), in **x** = (*x, y, z*) at time *t*.

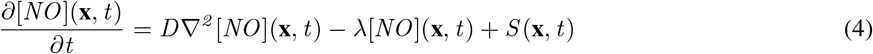

Where:

*D* is the diffusion coefficient. Thanks to low molecular weight and non-polarity, the NO can be considered to diffuse isotropically through the tissue, meaning that the diffusion coefficient is constant scalar. As reported in [Wood et al., 2011], we used a diffusion coefficient of 8.48 *×* 10^−10^ m^2^/s.

*λ*[*NO*](**x**, *t*) is the inactivation term. It represents a first-order reaction that governs the NO consumption in the brain tissue [Garthwaite, 2016], and it is defined by a rate constant of *λ* = 150 s^−1^, equivalent to a half-life for the NO molecule of 4.6 ms (values taken from [Wood et al., 2011]). This inactivation term can be seen as a simple global loss function that allows us to consider all background reactions involving the NO, i.e., with oxygen species and metals as well as with the *Heam* group of target sGC proteins.

*S*(**x**, *t*) is a function describing the dynamics and location of NO sources.

In order to solve Equation 4, we adopt some simplification on the geometry of the problem. First of all, we modelled each source of NO as a point source, from which the NO diffuses uniformly in all directions. We can safely assume radial symmetry to compute the diffusion profile in space. We compute [*NO*] with respect to the distance *r* from the source, not to **x** (3D Cartesian coordinates). This means that the source will have a fixed location in *r* = 0, and it will be described by just its evolution in time, hence *S*(*t*). To represent the action of multiple sources, we will simply sum each of their contributions with respect to a given point of observation. Moreover, due to the rapid decay in the NO concentration (high inactivation rate), we can assume a finite domain with boundaries condition being [*NO*](*r, t*) ≈ 0. We computed the solution of Equation 4 using Green’s function [Poirier and Poirier, 2016]. In our case, given:

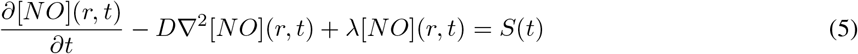

To summarize, we have developed a simulator that is capable of generating physiological concentration of NO upon neuronal activation through the nNOS sources and computing its spatial and temporal diffusion in all the pre-defined points of the surrounding volume.

### 2.2 NO-dependent plasticity model

To investigate the impact of the NO signal on cerebellar adaptation, we modified the spike-timing-dependent plasticity (STDP) learning rule developed to model LTP and LTD at *pf* -PC synapses [Antonietti et al., 2016, Garrido et al., 2013, Luque et al., 2016]. These modifications account for the involvement of NO in the plasticity process, aiming to improve the network’s adaptation in an eye-blinking classical conditioning protocol.

#### 2.2.1 Cerebellar inspired spiking neural network

First of all, we employed the cerebellar-inspired SNN developed in Geminiani et al. [2022], illustrated in Figure 1A. Each population is modelled with point neuron models. In particular, the mossy fibers (*mf* s) and the Glomeruli (Glom) are represented with *“parrot”* neurons, reporting in the output the input signal they receive. The Golgi Cells (GoCs), the Granule Cells (GrCs), the Stellate and Basket Cells (MLIs), and the PC are all represented with the Extended-Generalized Leaky Integrate and Fire (E-GLIF) model, which allows keeping the main electroresponsive features of neurons while reducing the computational load of simulations [Geminiani et al., 2018]. The Inferior Olivary Nuclei (IOs) are modelled as a conductance-based leaky integrate-and-fire neuron. The parameters (reported in Supplementary Table1) have been tuned for each population model in previous studies [Geminiani et al., 2022], as well as all the connection parameters of the network (Supplementary Table2). For the sake of simplicity, the neural connections are modelled with a static conductance-based synapse model, with delays extracted from literature and synaptic weights tuned to mirror the resting physiological firing rates observed in mice, in Geminiani et al. [2022]. Only the *pf* -PC synapses exert plasticity and follow an ad hoc STDP rule, where LTD is driven by the *cf* s teaching signal [Antonietti et al., 2016].

**Figure 1:**
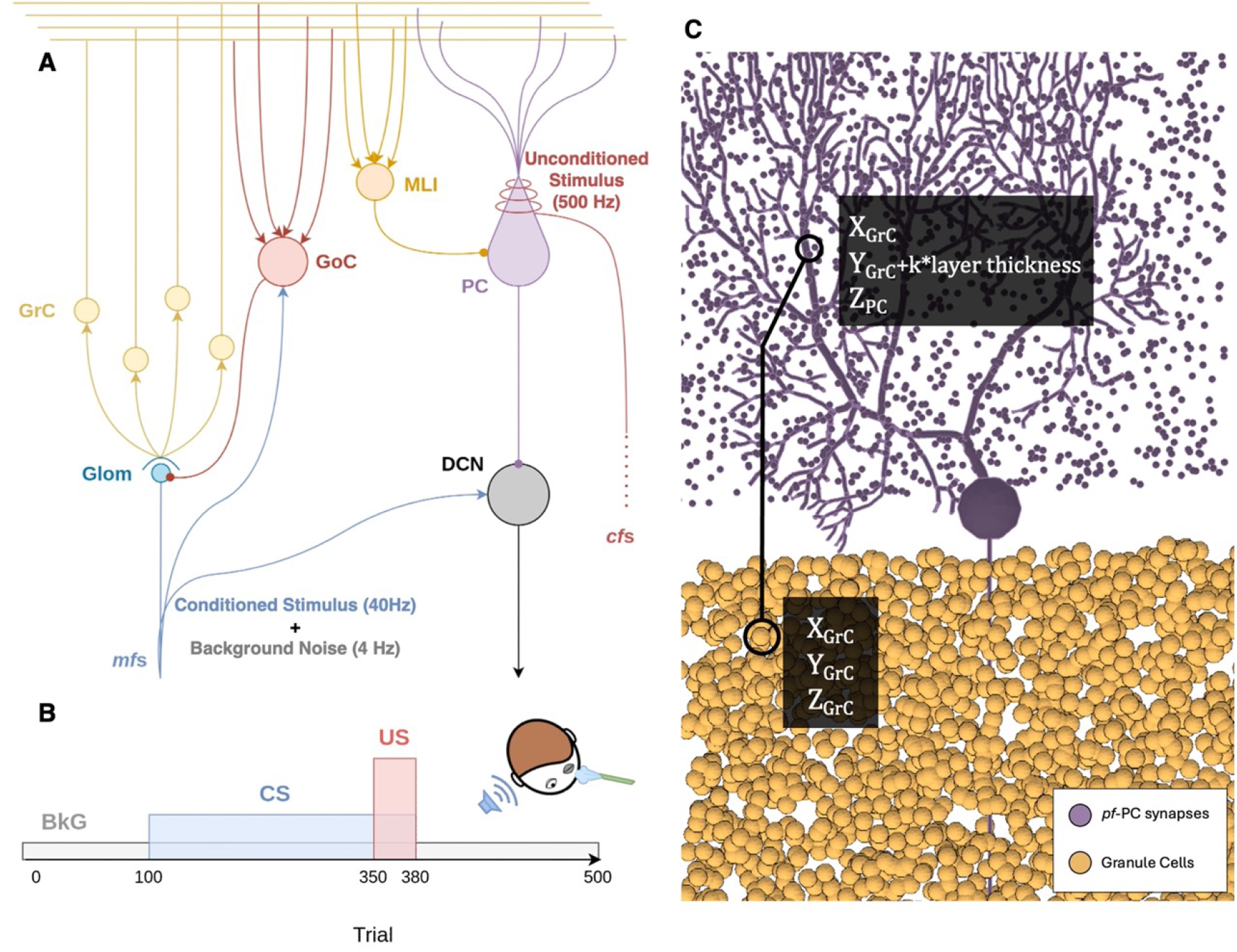
Cerebellar Microcircuit Model with EBCC protocol and nNOS placement: **A)** the network is composed by: mossy fibers (*mf* s), Glomerulus (Glom), Granule Cell (GrC), Golgi Cell (GoC), Stellate and Basket Cells (MLI), Purkinje Cells (PC), Inferior Olivary Nuclei (IO), climbing fibers (*cf* s), Deep Cerebellar Nucleus (DCN). **B)** Regarding the EBCC protocol A Conditioned Stimulus (CS) is conveyed by a subset of *mf* s, each receiving a non-recurrent 40 Hz spike train for 280 ms. The Unconditioned Stimulus (US) is a 500 Hz burst delivered to IOs for 30 ms. The CS and US stimuli co-terminate after 380 ms since each trial begins. The US causes a Conditioned Response (CR), i.e., an increase in the DCN firing rate. **C)** nNOS placement is based on the relative connected GrC (yellow) and PC (purple). The coordinates relations are: *x*_*nNOS*_ = *x*_*GrC*_, *y*_*nNOS*_ = *y*_*GrC*_ + *k* * *layer thickness, z*_*nNOS*_ = *z*_*P C*_

#### 2.2.2 nNOS placement on the Purkinje cell dendritic Tree

When simulating the NO signalling pathway, the geometrical organization of nNOS and the synapses in which NO is produced become crucial, as they determine the formation of the NO cloud and its influence on specific synapses. To reconstruct the geometrical arrangement of nNOS, we consider the synapse density on a single PC dendritic tree area. With an average of approximately 1500 active synapses and a mean dendritic tree area of 3 *×* 10^4^ *μm*^2^, we adopt a density of 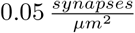 (values previously reported in De Schepper et al. [2022]). For precise placement of the parallel fiber *pf* -PC synapses, we adhere to the geometrical rule for *pf* -PC connections proposed by the Brain Scaffold Builder tool [De Schepper et al., 2022]. The synapse IDs are derived from the generated GrC-PC connection matrix. Each nNOS is assigned a geometrical point corresponding to a synaptic bouton on the respective PC dendritic tree, ensuring a unique association between each *pf* -PC synapse and its corresponding nNOS. The coordinates of the geometrical points associated with nNOS are determined as illustrated in Figure 1C. The *x* coordinate aligns with that of the corresponding GrC, as its ascending axon bifurcates perpendicularly in the Molecular Layer, crossing the PC dendritic tree. The *y* coordinate is proportional to the depth of the corresponding GrC in the granular layer. Finally, the *z* coordinate represents the position within the PC dendritic plane.

#### 2.2.3 Co-simulation NEST & NODS

To test the NO role as an enabler for plasticity, we run two sets of 10 simulations, 30 trials each, where only background noise is delivered to the *mf* input of the network. The results are then averaged for the 10 simulations. In the first 10 simulations, a standard STPD rule defines *pf* -PC weight updates in the cerebellar network. In the second 10 simulations, instead, the NO-dependent STDP rule controls *pf* -PC weight updates (Supplementary Table3). Then, in order to test the contribution of NO in the learning process of the cerebellum, we first reproduced the EBCC protocol proposed in Geminiani et al. [2022] for the physiological network, and we compared the network performance in the same conditions, but modifying the learning rule to account for NO diffusion. We used the spiking neural network simulator NEST [Gewaltig and Diesmann, 2007], a widely used framework that focuses on the dynamics, size and structure of neural systems. In the proposed EBCC protocol, a conditioned stimulus (CS) is conveyed by a subset of *mf* s, each receiving a non-recurrent 40 Hz spike train for 280 ms. The teaching signal, the unconditioned stimulus (US), is a 500 Hz burst delivered to IOs for 30 ms. The CS and US stimuli co-terminate after 380 ms since each trial begins (Figure 1B). All *mf* s receive a background noise modelled with a Poisson generator.

The synaptic weights of each *pf* -PC connection evolve following the STDP rule for LTD in and for LTP present in [Antonietti et al., 2016]. The plasticity rule is based on the observation that *pf* stimulation coupled with *cf* activation (teaching signal) triggers LTD at synapses between *pf* s and PCs receiving the *cf* signal, while *pf* stimulation alone causes LTP. We modified this learning rule to include NO-dependent plasticity. We first compute the NO concentration [*NO*] produced in each synapse, accounting also for the NO diffused from near synapses. In NEST, the device to transmit the teaching signal to the synapses is called *volume transmitter*. In the CerebNEST network’s original implementation, we find a single volume transmitter device for each PC, that does not have any physical properties and each connection would update its weight upon *pf* and *cf* activation in the same time window. When introducing the NO contribution, we have to take into account an additional constraint in the connections’ weight update: the relative distances of the synapses. In order to do so, we needed to assign spatial information to the synapses, thus to the *volume transmitter* devices, which in this case would be associated with each synapse of each PC. Because of this modification with respect to the original network, we had to re-tune the *A*_*plus*_ and *A*_*minus*_ in order to achieve the same learning curves as in Geminiani et al. [2022]. We used a grid searching algorithm to explore the combination of *A*_*plus*_ and *A*_*minus*_ (Supplementary Figure 1). As an index to determine the parameters to select, we used the difference in the firing rate of the PCs pre- and post-learning. By exploring the combination of *A*_*plus*_ and *A*_*minus*_ values around the one reported in Geminiani et al. [2022], we aimed at a depression of 20 Hz in the response of the PCs, evaluated in the interval where the conditioned response should be elicited, i.e., between 100 and 50 ms before the US. We found *A*_*plus*_ = 8 *×* 10^−5^ and *A*_*minus*_ = −4 *×* 10^−4^ to be the optimal learning rates.

To model the experimental findings regarding the plasticity enabler role of NO molecule (see subsection 1.3), we compute *G*_*NO*_, a sigmoid function that depends on the [*NO*] and ranges from 0 to 1 (see Equation 6), as an additional gain factor that modulates the learning rate *A*_*minus*_, in LTD equation, and *A*_*plus*_, in LTP one. By including *G*_*NO*_ we obtain the *“metaplasticity”* rule theorized by Schweighofer and Arbib [1998].

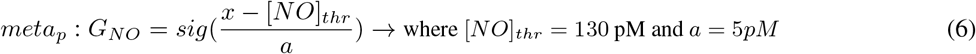

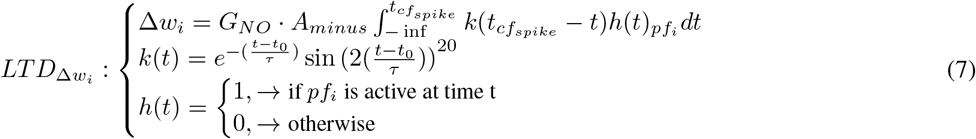

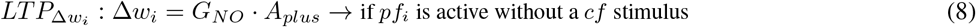

#### 2.2.4 Compare standard and NO-dependent STDP

To test the hypothesis formulated in many experimental studies [Lev-Ram et al., 2002, Maffei et al., 2003, Garthwaite, 2016], that the NO enables plasticity when relevant signals are delivered to the network, we firstly simulate the case scenario where only background noise is present. In particular, we evaluate the rate of weight’s updates, computed as the mean number of updates for each synapse upon different background noise levels fed to the network. Instead, to evaluate how the learning changes upon the introduction of NO dependency we evaluate the PC firing modulation during complete EBCC simulations. In particular, we compare the EBCC featuring the standard STDP at three different background noise levels (0 Hz, 4 Hz and 8 Hz) with the respective simulations that featured NO-dependent STDP.

To assess firing modulation, we computed the Spike Density Function (SDF) as the convolution of each cell’s spikes in every trial, with a Gaussian kernel of 20 ms for PCs [Dayan and Abbott, 2001]. SDFs of the population were then computed by averaging SDFs of individual cells. To quantify learning in terms of neural activity, an index of SDF change was computed in the expected CR time window, between 250 ms and 300 ms, and over the trial baseline, between 150 ms and 200 ms from the CS onset. The SDF change index is computed for each trial as the SDF mean in the time window of interest subtracted by the SDF mean in the same time window of the first trial. This quantifies the time-locked rate change, i.e., the modulation in the inter-stimuli interval within each trial.

## 3 Results

### 3.1 Single source model

To validate the system of differential equations describing the synthesis process we compared the results with the ones obtained in a different simulation environment, NEURON [Kumbhar et al., 2019]. This simulator allows the building of detailed individual neuron properties using compartmental models. In particular, we exploited its ability to simulate, with a high level of realism, the biochemical reactions taking place in the intracellular space using the Reaction&Diffusion module. The biochemical cascade resulting in the NO synthesis has been triggered in both models by two 40 Hz burst stimuli 100 ms long, with an inter-burst interval of 300 ms. In Figure 2 are shown the spiking activity of neurons (Figure 2A) and the relative intermediate compounds concentration along time, leading to NO production (Figure 2C). The comparison between the two simulation environments expresses an almost perfect superimposition of the results. This permits the validation of the NODS model with the widely used NEURON simulation platform. Furthermore, the results show that the lower computational load production function of NODS is able to replicate the results, by depending on the time of spike events. Thus, the implementation is compatible with information encoded in a spiking neural network: binary time series. Once we have obtained a reliable description of the production function, we simulated the diffusion from a single source. Figure 3 shows the time and space profile of the NO signal, produced by a single source stimulated with different stimulation conditions. The curves show how different stimulation frequencies result in a different distance covered by the NO signal. In particular, the higher the frequency, the more NO is produced, and the larger area is covered by the NO signal (Figure 3B). However, the model reaches a physiological saturation point in the amount of NO that can be produced and consequently diffused between 100 Hz and 300 Hz. The evaluation of the maximal amount of NO that can be produced by a single source, around 130 pM, serves us to set a threshold to the amount of NO that would enable plasticity, in Equation 6. We also compared the diffusion from a single source simulated with our approach with the results of Wood et al. [2011], where an empirical approach has been used. In Figure 4 we compared the two models ^2^ by evaluating the NO concentration at a distance of 200 nm from the source following a single spike activation. As result we obtain a clear overlap of NODS simulator diffusion curve of a single source with the Garthwaite results Garthwaite [2016].

**Figure 2:**
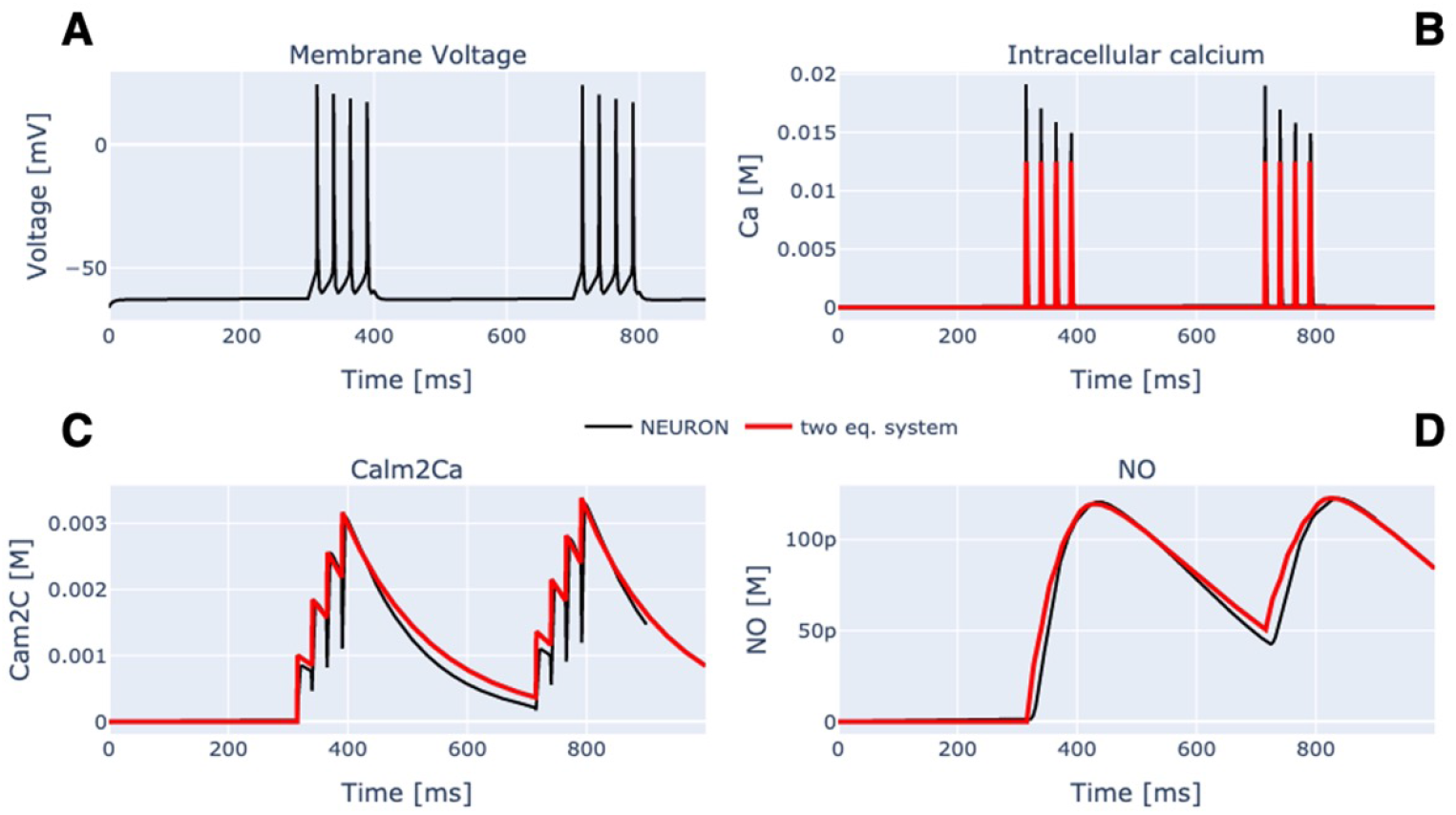
Compare NODS and NEURON Reaction&Diffusion module: Black curves refer to results obtained using the NEURON simulation environment, while red curves to NODS.

**Figure 3:**
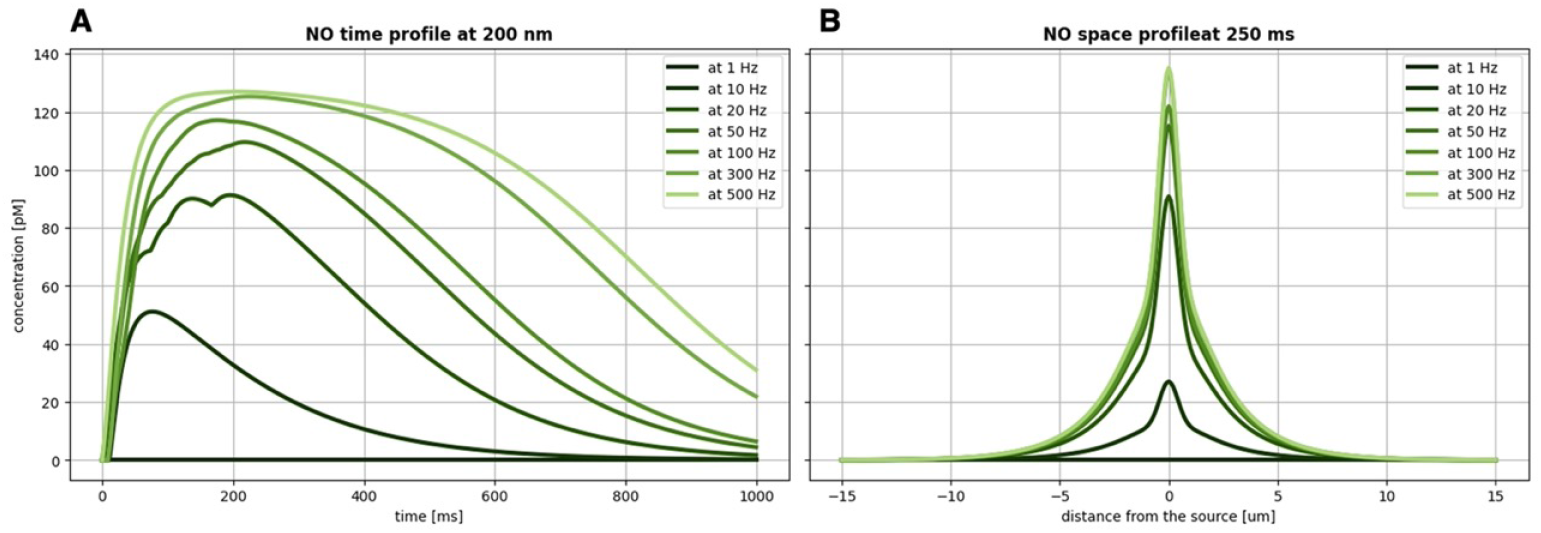
NO produced and diffused from a single source: time **(A)** and space **(B)** profile of the NO signal, produced by a single source stimulated at with different intensities (colour-coded by darkening the shade of green): single spike, 10 Hz, 20 Hz, 50 Hz, 100 Hz, 300 Hz, 500 Hz. All stimuli have been delivered at *t*_*i*_ = 0 ms and last for 200 ms.

**Figure 4:**
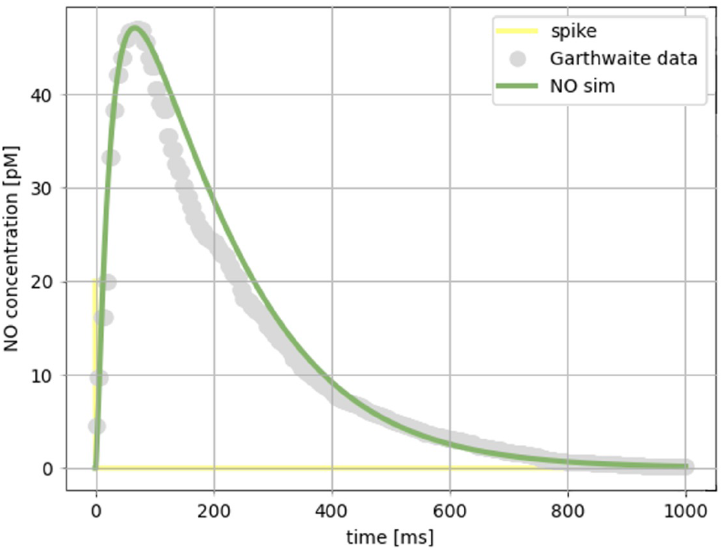
Comparison with Garthwaite [2016]: NO concentration at a distance of 200 nm from the source following a single spike activation.

### 3.2 Only background noise simulation

As reported in both modelling and experimental studies [Lev-Ram et al., 2002, Garthwaite, 2016, Hardingham et al., 2013, Ogasawara et al., 2007] only neighbouring small groups of *pf* s activated close in time at a certain frequency would exert plasticity or, in simulation terminology, update their (synaptic) weights, thanks to the presence of a NO cloud produced in the area where each group is located. From the results of the 10 only background noise simulations, with and without the NO-dependent plasticity, we observed that with the NO enabling mechanism, each weight is less likely to update. In Supplementary Table3 we report the rate of weight updates for one simulation with three different noise levels: 0 Hz, 4 Hz, and 8 Hz. The rate of weight updates are calculated by taking all the weight updates for each synapse and dividing them by the total simulation time. We then averaged for all the synapses and all the simulations. Given the hypothesis that the background noise does not correlate in a specific space of the granular layer, it activates “isolated” *pf* s, resulting in smaller NO clouds not capable of influencing neighbouring synapses. Thus, these synapses will be less likely to reach the threshold value of NO required to update.

### 3.3 NO-dependent plasticity in an EBCC protocol

In Figure 5, we compared the PCs response as SDF amplitude (averaged for the 10 simulations) for 30 trials where the network features the standard STDP during an EBCC protocol vs. a network that receives the same stimuli but features a NO-dependent STDP rule.

**Figure 5:**
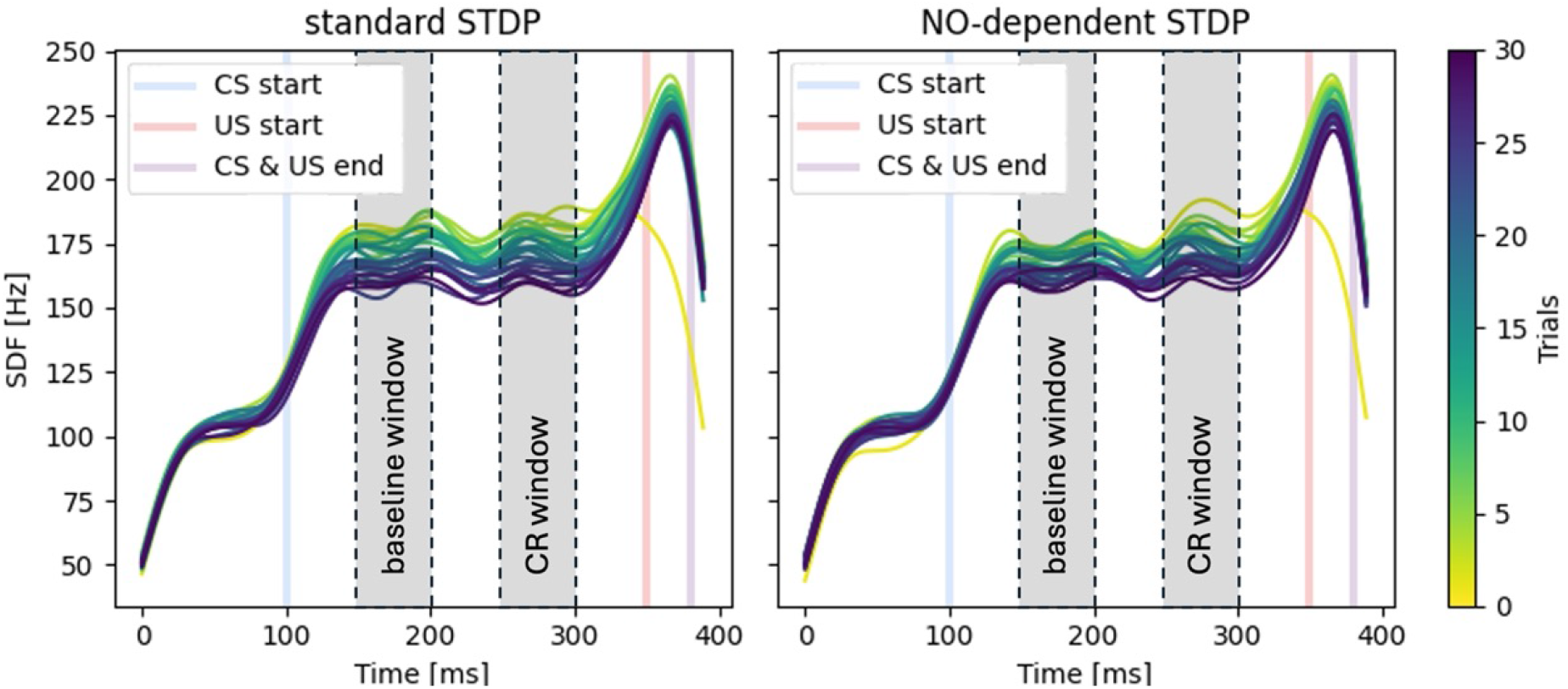
Comparison of PC SDF over trials with standard STDP and NO-dependent STDP: both panels report the SDF in PC population (averaging across cell SDFs) for each trial. The progress of the trials is colour-mapped by the vertical colour bar on the right. The vertical lines represent the CS-onset (light blue), US-onset (light red), and CS-US co-termination (lilac). In grey are shown the two windows, baseline and CR window, for evaluation of functional plasticity. On the left panel, the results of EBCC simulated using the standard STDP rule. On the right panel, the results of EBCC simulated using the NO-dependent STDP rule.

To test the hypothesis that the NO facilitates the selection and updating of synapses according to the relevance of a given context, intrinsically encoded in the proximity of the given synapses, we investigate whether the presence of NO would increase the robustness of the learning efficiency to different levels of background noise. Figure 6A shows the trend over trials of the SDF change computed in the conditioned response window and in the baseline window, highlighted in Figure 5. We can see that in the EBCC simulation where the NO-dependent STDP governs learning, the PC depression is more pronounced in the CR window with respect to the baseline window. Thus, the latter is less likely to provoke an early CR right after the CS onset. In the case with NO, the time-locked anticipatory action is more precise in time, being confined to happen just before the US delivery. What we expect from an efficient learning system tested with an EBCC protocol is to have a greater SDF change (depression) during the CRwindow with respect to the baseline window. According to the STDP rule, the LTD should be stronger between 50 ms and 100 ms before the US stimulus, to elicit a correct anticipatory response. Using a standard STDP rule, we notice that this is true for the last 5 trials, where the baseline SDF change starts to diverge from the CR-window SDF change in the simulated conditions with 4 Hz (black curves in Figure 6A) background noise. The trends of the CR-window and baseline results are similar respectively between the standard STDP and NO-STDP implementation with 0 Hz background noise. In the condition of 8 Hz background noise, we cannot distinguish a separation between the CR-window SDF change curve and the baseline SDF change curve in both conditions. With a NO-dependent STDP rule, we observe that with a 4 Hz (green curves in Figure 6A) background noise level, the two SDF change curves, the one computed for the baseline and the one computed for the CR-window, remains quite distant since the beginning, meaning that the baseline interval is less changed by the *pf* -PC plasticity, as it should. In particular, we can see that the baseline SDF change is more stable over the trials and smaller than the baseline SDF change for the standard STDP rule.

**Figure 6:**
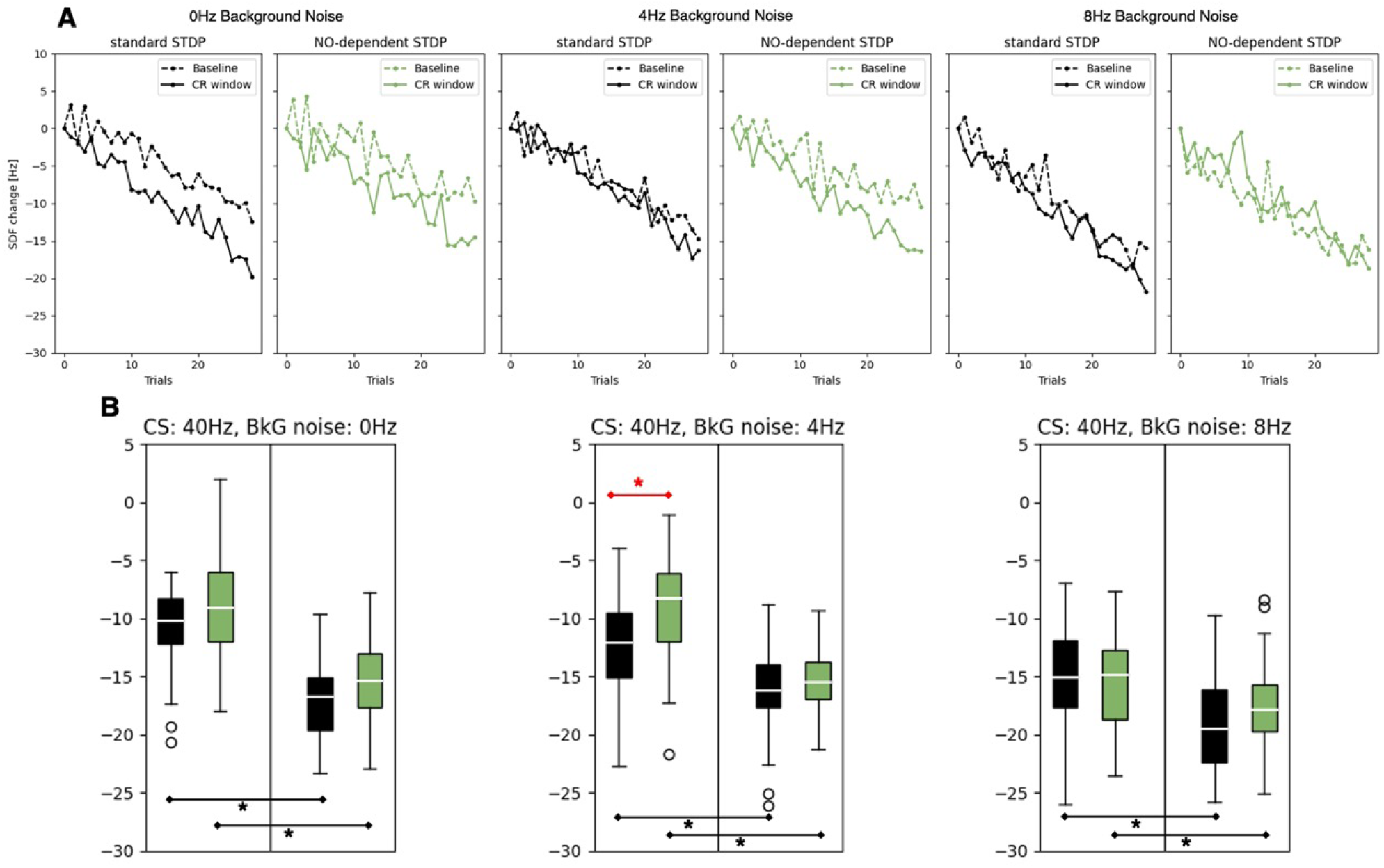
SDF change for different levels of background noise: Standard STDP is colour-coded in black and NO-dependent STDP is colour-coded in green. **A)** Dashed lines connect SDF change computed in the baseline window in each trial, while continuous lines connect SDF change computed in the CR response window. **B)** SDF change in PC from the last 5 trials, for EBCC protocols with a background noise of 0 Hz (left), 4 Hz (center), and 8 Hz (right). Standard STDP is colour-coded in black and NO-dependent STDP is colour-coded in green. * corresponds to p < 0.01 in the Wilcoxon test.

In Figure 6B, we compare the SDF change in the last 5 trials, averaging over 10 simulations, and evaluate the statistical differences between the standard STDP and the NO-dependent STDP EBCC performance, by using a Wilcoxon statistical test with statistical significance at p < 0.01.

With 4 Hz background noise, the baseline SDF changes for the NO-dependent STDP (green boxplots) is significantly smaller than the baseline SDF changes for the standard STD case (black boxplot), with a *p* − *value* = 5.31 *×* 10^−9^. This means that a NO-dependent STDP rule impacts the learning efficiency by better discriminating the actual meaningful interval to be learned (CR-window) from the baseline. Moreover, it is interesting to notice that in the case of 0 Hz background noise with NO-dependent STDP rule, the SDF changes, in both intervals, are not significantly different from their counterpart in the standard STDP simulation. On the other hand, for 4 Hz background noise, the CR-window SDF change is significantly greater when simulated with the NO-dependent STDP rule than in the standard STDP simulation, granting faster learning. However, when we inject a higher level of noise in the network input, i.e., 8 Hz, the NO-dependent STDP performs similarly to the standard STDP rule (no significant differences).

## 4 Discussions

We began this work from the guesses about NO physiological role in the cerebellum and we chose to focus at the *pf* -PC synapse level for two main reasons: the well-known functional role of PC cells in cortico-cerebellar microcircuits, which collect all the inputs in its dendrites and produce the sole output of the circuit Albus [1971], Nguyen-Vu et al. [2013]; the geometrical spacing properties of the PC dendritic trees and the *pf* -PC synapses, where NO fits well with its possible functional role associated to metaplasticity. Since we were interested in how the NO signal influences the cerebellar adaptation, we modified the STDP learning rule for *pf* -PC synapses to incorporate the dependency of NO in the plasticity mechanism, trying to replicate experimental findings [Lev-Ram et al., 2002, 1997, Wang et al., 2014]. The simulations and models are validated through testing during an EBCC protocol to understand the significance of NO in cerebellar learning. In this context the presented results bridge the gap between molecular processes at single synapses and network-level learning mechanisms, showcasing how NO diffusion translates into functional outcomes in cerebellar plasticity.

### 4.1 Single source model and the NODS simulator

The development of the production and diffusion model for a single NO source is an important starting point for understanding the role that NO plays in cerebellar plasticity. NODS model tries to replicate the biochemical and physical processes underlying NO diffusion and provides a critical link between molecular activities and synaptic changes at the network level. By simulating the spatiotemporal dynamics of NO diffusion from a single nNOS enzyme source, the model captures the balance between NO production rates, its diffusion through the neural tissue, and the eventual inactivation or consumption of NO molecules. NODS is developed to work in an SNN framework. Thus, it has to ensure a low computational load, so as to not burden the simulation time of large-scale networks, and shape NO biochemical cascade and kinetics dependently on spike trains. Although these simplifications, our NO production algorithm gives results in line with the ones obtained with NEURON, which, however, are based on multi-compartmental models. Altogether this testifies to both NODS validation and its improvement with respect to state-of-the art simulators. Furthermore, the biochemical cascade initiated by the activation of nNOS leads to NO production and its diffusion profile. This is characterized by a rapid decay and a constrained spatial distribution, in line with [Garthwaite, 2016] production and diffusion model, which underpins the localized yet significant impact of NO on synaptic plasticity. From the point of view of NO functionality, the fine-tuned regulation ensures that NO-mediated signalling remains confined but effective, influencing synaptic efficacy and learning rates within a delimited spatial domain.

### 4.2 Role of NO in the cerebellar circuitry

By understanding the initial steps of NO signalling at the level of individual sources, we gain a deeper appreciation for the complex interplay between molecular dynamics and network-level learning mechanisms. The simulations underscore NO’s capacity to fine-tune synaptic weights, dynamically adjusting the balance between LTP and LTD based on the temporal correlation of synaptic inputs with error signals. This dynamic adjustment, a hallmark of metaplasticity, is crucial for the cerebellum to prioritize learning from synaptic inputs most relevant to executing accurate motor commands. Through modelling and the simulation of NO-dependent STDP in the cerebellar circuitry, we tried to test the hypothesis formulated by Schweighofer and Arbib [1998] on how NO influences cerebellar learning and plasticity. Their model relies upon the findings that nonspecific long-term depression, which we could associate with NO-dependent plasticity, plays a crucial role in enhancing the resilience of a modelled PC against noise arising from the specific activation of synapses. If this spatial noise or variability in synaptic activity represents natural errors or fluctuations in sensory signals or motor commands, nonspecific plasticity could potentially serve as a mechanism for error correction, pattern generalization, and completion. With this in mind, we wanted to test NO influence on a cortico-cerebellar SNN activity, focusing on three pivotal aspects: control learning rates of parallel fiber-Purkinje synapses, addressing the overfitting problem, and the relevance of inputs alongside learning interference.

#### Learning rates of *pf* -PC synapses

The simulation results underscore the critical role of NO in dynamically regulating the learning rates of parallel fiber-Purkinje cell synapses. By modulating NO production and diffusion in response to synaptic activity, our model demonstrates a mechanism through which the cerebellum can adjust learning rates, enhancing its capacity for rapid adaptation to new motor tasks or environmental changes. This dynamic adjustment mechanism, underpinned by NO signalling, aligns with the theoretical considerations for optimal learning rates, ensuring that synaptic modifications occur at a pace conducive to stable and efficient learning without succumbing to oscillations or saturation.

#### Addressing the overfitting problem

Our findings also contribute to understanding how the cerebellum addresses the potential issue of overfitting, given the vast number of synapses per Purkinje cell. The selective mechanism is influenced by the spatial and temporal dynamics of NO diffusion. From a computational point of view NO concentration increases following repetitive activity in neighbouring synapses which in turn activates plasticity only in those ones included in a cloud of [*NO*] above the threshold. On the other hand, from a functional point of view, this acts through a selection of inputs coming from *pf* s. This suggests a natural way the cerebellum enhances its generalization power, preventing overfitting by filtering out noise and irrelevant information from the learning process. The NO-dependent modulation of synaptic plasticity, by promoting a selective and context-dependent adjustment of synaptic efficacy, plays a crucial role in focusing learning on relevant synapses while minimizing adjustments to those less involved in the task at hand.

#### Relevance of inputs and learning interference

The simulations further illustrate how NO-mediated plasticity mechanisms could mitigate learning interference by differentiating between relevant and irrelevant inputs. NO diffusion serves as a gating mechanism: it associates higher learning rates with synapses experiencing temporally correlated activity with error signals and reduces the influence of synapses with less relevant input. This comes directly from the NODS implementation and NO kinetics properties. Due to the low concentrations at which NO is present, its signalling range is often considered limited to a short distance around its sources. However, NO’s influence extends beyond a strictly localized scope. Coordinated nNOS activity among a population of neurons leads to an accumulation of NO concentration in the extracellular space, reaching levels capable of eliciting neuronal stimulation in that vicinity. This selective enhancement, or suppression, of synaptic changes, is in line with the results presented by the [Ogasawara et al., 2007] model. In our study, applying it during a motor learning functional protocol, we can assess that NO plasticity contributes meaningfully to motor command learning, thereby reducing the potential for interference and enhancing the cerebellum’s ability to construct accurate internal models.

## 5 Concluding remarks

This project aimed at bridging the gap between molecular processes and network-level learning mechanisms, showcasing how NO diffusion translates into functional outcomes in cerebellar plasticity. Leveraging advanced computational models, we simulate NO diffusion across a large-scale, physiologically inspired cerebellar network. This endeavour not only includes NO’s pivotal role as a modulator of synaptic efficacy but also its function in dynamically adjusting the cerebellum’s learning capabilities through the lens of metaplasticity. By simulating varying levels of NO diffusion, we demonstrated that higher NO concentrations, typically resulting from concerted synaptic activity, lead to a more pronounced adjustment in the synaptic learning rates. This phenomenon aligns with the theoretical framework of metaplasticity, where the cerebellum’s ability to adapt its learning rates based on the relevance and timing of inputs is essential for refining motor control and learning new motor tasks efficiently. NO-mediated plasticity allows the cerebellum to implement a more sophisticated strategy for learning and updating motor commands while reducing learning interference from irrelevant synaptic inputs. By modulating the learning rates through NO signalling, the cerebellum can effectively filter out, up to a certain degree, “noise” from synaptic inputs not directly involved in the current motor task, thus preserving the fidelity of motor command learning. This strategy is especially relevant in the context of complex motor tasks requiring the integration of kinematic information from multiple joints. Notably, we develop an efficient implementation of NO diffusion, which not only facilitates our large-scale simulations but also provides a tool for the scientific community to interface with different types of simulation tools. This enables the investigation of NO diffusion in various scenarios, including different brain areas at different scales or even combining the simulation with artificial neural networks. Additionally, the developed methodologies and tools have the potential to contribute to future studies investigating NO dynamics and its interactions with neural activity and neurovascular coupling in various brain regions.

### 5.1 Limitations and future developments

The current implementation of NODS is a flexible Python module capable of simulating NO diffusion across different network configurations. It effectively models scenarios where the spiking activity stimulates nNOS, given the coordinates of nNOS and the evaluation points for NO concentration. However, NODS implementation could be significantly improved, in terms of simulation times, by parallelizing the computation of the diffused NO from each nNOS source. Future enhancements could focus on improving NODS compatibility with simulators that do not rely exclusively on point neuron models. For example, by integrating inputs of actual *Ca*^2+^ concentrations that trigger NO production in compartments where nNOS is located, NODS can provide more efficient simulations of NO dynamics in complex neuronal architectures. Furthermore, given the presence of nNOS also in the granular layer and the role of NO in plasticity, it would be significant to implement the NO-dependent plasticity mechanism also at the *mf* s-GrCs synapse level. Another possible development path involves integrating NODS into larger simulations that include representations of the vascular system. This would enable the study of neurovascular coupling phenomena, necessitating the inclusion of endothelial NO synthase sources. Modifying NODS to account for endothelial NO synthase parameters will be crucial for accurately modelling NO production and diffusion in contexts where blood flow and vascular responses play significant roles in neural activity and plasticity. By pursuing these enhancements, NODS can become an even more powerful tool for researchers, facilitating comprehensive studies that span from intracellular processes to system-wide interactions.

## Acknowledgements

The work of AA, AP, BG, and CAS in this research is also supported by Horizon Europe Program for Research and Innovation under Grant Agreement No. 101147319 (EBRAINS 2.0).

The work of AMT was supported by a Voucher (CEoI 4-Rodent microcircuits: RisingNet Whole-bRaIn rodent SpikING neural NETworks) from the European Union’s Horizon 2020 Framework Programme for Research and Innovation under the Specific Grant Agreement No. 945539 (Human Brain Project SGA3). The simulations in NEURON were implemented by Stefano Masoli, Department of Brain and Behavioral Sciences, Universitá di Pavia, Pavia, Italy.

## Footnotes

1 A spike train stimulates NMDA receptors. Ca^2+^ enters the intracellular space through the NMDA receptor, increasing the intracellular concentration of Ca^2+^. Ca^2+^ react with calmodulin, and the concentration of Calm2C (calmodulin/calcium bounded) increases. This induces a catalytic activation of the nNOS enzyme that starts producing NO using oxygen and reducing NADPH to catalyze the conversion of arginine to citrulline.

2 The corresponding author of [Wood et al., 2011] kindly provided annotated Mathcad worksheets to reproduce their results

