## Supplementary Material for "Modeling Nitric Oxide Diffusion and Plasticity Modulation in Cerebellar Learning"

July 24, 2024

| ***EGLIF model*** | **Granule** | **Golgi** | **Purkinje** | **Basket** | **Stellate** |
|---|---|---|---|---|---|
| $C_m$ $[pF]$ | 7 | 145 | 334 | 14.6 | 14.6 |
| $E_L$ $[mV]$ | -62 | -62 | -59 | -68 | -68 |
| $I_e$ $[pA]$ | -0.888 | 16.214 | 742.543 | 3.711 | 3.711 |
| $V_{reset}$ $[mV]$ | -70 | -75 | -69 | -78 | -78 |
| $V_{th}$ $[mV]$ | -41 | -55 | -43 | -53 | -53 |
| $t_{ref}$ $[ms]$ | 1.5 | 2 | 0.5 | 1.59 | 1.59 |
| $A_1$ $[pA]$ | 0.01 | 259.988 | 157.622 | 5.953 | 5.953 |
| $A_2$ $[pA]$ | -0.94 | 178.01 | 172.622 | 5.863 | 5.863 |
| $E_{rev1}$ $[mV]$ | 0 | 0 | 0 | 0 | 0 |
| $E_{rev2}$ $[mV]$ | -80 | -80 | -80 | -80 | -80 |
| $E_{rev3}$ $[mV]$ | 0 | 0 | 0 | 0 | 0 |
| $E_{rev4}$ $[mV]$ | N/A | -80 | N/A | N/A | N/A |
| $V_{init}$ $[mV]$ | -62 | -62 | -59 | -68 | -68 |
| $V_{min}$ $[mV]$ | -150 | -150 | -350 | N/A | N/A |
| $K_1$ $[ms^{-1}]$ | 0.311 | 0.031 | 0.195 | 1.887 | 1.887 |
| $K_2$ $[ms^{-1}]$ | 0.041 | 0.023 | 0.041 | 1.096 | 1.096 |
| $k_{adap}$ $[MH^{-1}]$ | 0.022 | 0.217 | 1.492 | 2.025 | 2.025 |
| $lambda_0$ $[adim.]$ | 1 | 1 | 4 | 1.8 | 1.8 |
| $tau_V$ $[ms^{-1}]$ | 0.3 | 0.4 | 3.5 | 1.1 | 1.1 |
| $tau_m$ $[ms^{-1}]$ | 24.15 | 44 | 47 | 9.125 | 9.125 |
| $tau_{syn1}$ $[ms^{-1}]$ | 1.9 | 0.23 | 1.1 | 0.64 | 0.64 |
| $tau_{syn2}$ $[ms^{-1}]$ | 4.5 | 3.3 | 2.8 | 2 | 2 |
| $tau_{syn3}$ $[ms^{-1}]$ | N/A | 0.5 | 0.4 | 1.2 | 1.2 |
| $tau_{syn4}$ $[ms^{-1}]$ | N/A | 2.4 | N/A | N/A | N/A |

| ***LIF model*** | **IO Nuclei** |
|---|---|
| $C_m$ $[pF]$ | 189 |
| $E_L$ $[mV]$ | -45 |
| $I_e$ $[pA]$ | 0 |
| $V_{reset}$ $[mV]$ | -45 |
| $V_{th}$ $[mV]$ | -35 |
| $t_{ref}$ $[ms]$ | 1 |
| $g_L$ $[nS]$ | 17.18 |
| $tau_{syn_{ex}}$ $[ms^{-1}]$ | 1 |
| $tau_{syn_{in}}$ $[ms^{-1}]$ | 60 |

Table 1: **Neuron model parameters** from Geminiani et al. [2022]

| **Connection Model** | ***Weight [nS]*** | ***Delay [ms]*** |
|---|---|---|
| **parallel fiber to purkinje** | 0.4000 | 5 |
| **mossy to glomerulus** | 1.0000 | 1 |
| **glomerulus to granule** | 0.2322 | 1 |
| **golgi to granule** | 0.2414 | 2 |
| **glomerulus to golgi** | 0.2402 | 1 |
| **golgi to golgi** | 0.0070 | 4 |
| **ascending axon to golgi** | 0.8228 | 2 |
| **parallel fiber to golgi** | 0.0538 | 5 |
| **parallel fiber to basket** | 0.1003 | 5 |
| **parallel fiber to stellate** | 0.1780 | 5 |
| **stellate to purkinje** | 1.6417 | 5 |
| **basket to purkinje** | 0.4357 | 4 |
| **stellate to stellate** | 0.0046 | 4 |
| **basket to basket** | 0.0058 | 4 |
| **ascending axon to purkinje** | 0.8820 | 2 |
| **io to purkinje** | 300.00 | 4 |

Table 2: **Synaptic connection parameters** from Geminiani et al. [2022]

| ***Noise rate*** | ***Standard STDP [num. weight updates/ms]*** | ***NO-dependent STDP [num. weight updates/ms]*** |
|---|---|---|
| 0 Hz | 0.50698 | 0.31139 |
| 4 Hz | 0.50701 | 0.31256 |
| 8 Hz | 1.02817 | 0.70240 |

Table 3: **Rate of weight's updates in each *pf*-PC synapses on different noise level** comparison of the rate of weight updates for one simulation with three different noise levels (0 Hz, 4 Hz, and 8 Hz) between standard and NO-dependent STDP .

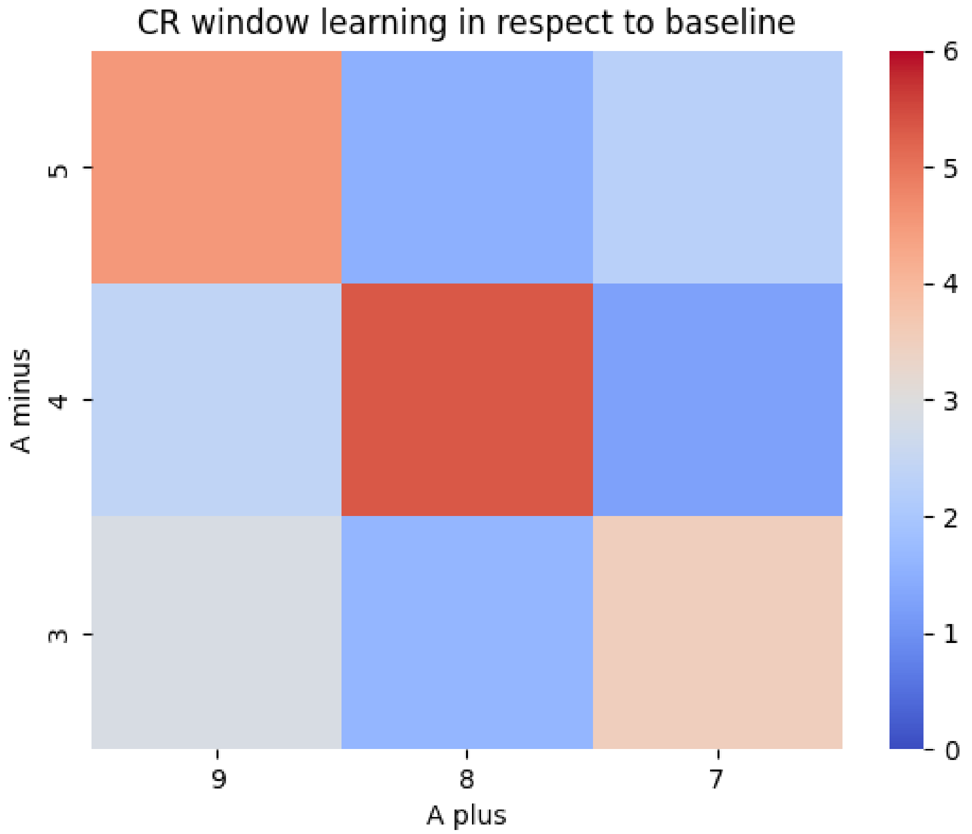


Figure 1: **Grid search for $A_{plus}$ and $A_{minus}$:** the colourmap reports the matrix of the combinations for $A_{plus}$ and $A_{minus}$. For every combination, the average difference of SDF change for the last 5 trials for the baseline window and the CR window is shown. Both the SDF changes are negative values, so the difference should be positive, in order to have a higher learning in the CR window with respect to the baseline. In the colourmap, the colour coding is red for positive values and blue for negative ones. Within the grid search, we seek the highest value of our metrics, which corresponds to the $A_{plus}$ and $A_{minus}$ to maximize learning.
